# A temporary challenge by tumor cells can lead to a permanent partial-impairment of memory CD8 T cell function

**DOI:** 10.1101/2025.02.17.638652

**Authors:** Daphné Laubreton, Margaux Prieux, Sophia Djebali, Maxence Dubois, Simon De Bernard, Olivier Gandrillon, Christophe Arpin, Jacqueline Marvel

## Abstract

Memory CD8 T cells typically exhibit improved effector functions compared to their naive counterparts. However, under certain activation conditions, such as chronic viral infections or cancer, these cells may develop functional defects. In this study, we compared the functional quality of memory CD8 T cells generated following tumor rejection with those arising from an acute viral infection. We found that tumor-induced (Tum-CD8) memory cells exhibited a distinct phenotype and transcriptomic profile compared with viral-induced (Vir-CD8) memory cells. These memory cells are characterized by the expression of inhibitory receptors and displayed altered functions including reduced IFNγ and TNFα production as well as changes in integrin expression. Additionally, the protective capacity of Tum-CD8 memory cells was flawed relative to that of Vir-CD8 memory cells. Importantly, the functional defects of Tum-CD8 persisted upon viral recall. Together, these findings indicate that transient tumoral stimulation can imprint a stable partial exhaustion-like program on memory CD8 T cells.

## Introduction

Following an acute infection, naive CD8 T cells undergo extensive proliferation and differentiate into cytotoxic effector cells that control and clear the infection. After the resolution of the inflammation, most effector cells die, whereas a small fraction differentiates into memory cells. Unlike their naive counterparts, memory cells persist for extended periods (1–3) and exhibit enhanced responsiveness upon antigen reencounter (4,5). This state of hyper-responsiveness results from transcriptional and epigenetic modifications established upon activation and maintained thereafter (6–8). As a result, the functional properties of memory CD8 T cells are modified leading to immediate proliferation and rapid acquisition of effector functions, such as cytotoxicity or cytokine production, following recall. Moreover, they adopt new trafficking and homing potentials (9–14).

In contrast, under conditions of persistent antigen exposure, such as chronic infections or cancer, memory CD8 T cells enter a state of hyporesponsiveness, referred to as exhaustion (15). This state is characterized by impaired cellular functions, including reduced cytokine production, diminished proliferative capacity, and decreased cytotoxicity (15). Exhausted CD8 T cells are characterized by a sustained expression of multiple cell-surface inhibitory receptors, including PD1 or TIM-3 (15–18) as well as distinct transcriptomic and epigenetic programs (18–22), in both chronic infection or tumorigenesis. These cells originate from a precursor population (Tpex) that is characterized by high TCF1 expression (23–26).

While exhausted CD8 T cells arising in chronic infections and cancer share common exhaustion features, they also exhibit specific characteristics (18,22,25). In an antiviral environment, Tpex are generated in both acute or chronic context upon strong TCR priming and require continuous antigen exposure for their maintenance (27,28). However, in cancer, T cell priming is inefficient because tumor antigens are not presented in a proper inflammatory and co-stimulatory context, and that can lead to a state of anergy (18,29). Indeed, CD8 T cells display altered cytokines production within the first few hours following *in vivo* activation by a tumor, a change that is associated with epigenetic modifications (30,31). These modifications include increased accessibility of the Pdcd1 gene-enhancer, which regulates PD-1 expression (30,31). This state of unresponsiveness can lead to exhaustion, if the tumor growths, leading to chronic stimulation of tumor-antigen specific T cells. However, the impact of a lack of effective costimulatory signals, in the absence of chronic antigen-exposure, on the responsiveness capacities of memory CD8 T cells remains to be fully elucidated.

To answer that question, we conducted a detailed comparison of the functional properties of memory CD8 T cells generated following tumor rejection, using acute viral stimulation as a gold standard for the generation of functional memory CD8 T cells. Both the virus (vaccinia virus - VV) and the tumor (EL4 cells) expressed the NP68 epitope recognized by the F5 TCR transgene, and are able to induce the activation and proliferation of F5-transgenic CD8 T cells. In these settings, the virus was cleared or the tumor was rejected within 2 weeks of exposure. We showed that memory CD8 T cells generated following the tumoral challenge displayed a distinct phenotype characterized by decreased integrins expression and increased expression of the inhibitory receptors PD1 and TIM-3. This exhausted-like phenotype is associated with a reduced capacity for cytokines production and proliferation upon re-activation leading to a defective protective function. Finally, we demonstrated that this phenotype was stable and could not be reversed, even after a subsequent viral challenge. Thus, our findings suggest that the suboptimal co-stimulation associated with the tumoral challenge is sufficient to imprint a partial exhausted-profile on memory CD8 T cells.

## Results

### Tumor or virus-induced memory cells display a restricted number of phenotypic and transcriptional profile differences

To determine the impact of a transient tumoral challenge on the generation of CD8 memory cells, we compared the phenotype of memory CD8 T cells generated after a viral challenge with VV or a tumor challenge with EL4 cells. Maximal viral replication was detected 5 days post-challenge (dpc) while tumor size peaked between 6 and 7 dpc (Fig.1a). The virus and tumor were both eliminated from the challenge site within 7 and 12 dpc respectively (Fig.1a). We first followed the number of F5 CD8 T cells over time in the blood and showed that the number of circulating cells peaked at 11 dpc for both virus-challenged or tumor-challenged mice (Fig.1b). Furthermore, the numbers of CD8 T cells recovered in the memory phase after a viral or a tumoral challenge were similar (Fig.1b). We then used Ki67 and Bcl2 labelling to compare the generation of early effector (EE, Ki67+ Bcl2-), late effectors (LE, Ki67- Bcl2-) and memory cells (Ki67- Bcl2+)(32). The kinetics of EE generation, as well as the expression of Ki67 by CD8 T cells, were similar between virus- and tumor-challenged mice (Fig.1c, Supplementary Fig.1a-b). However, the numbers of LE were maintained for a longer period of 3 to 5 days, and there was similar time delay in the appearance of memory cells following tumor challenge compared to viral challenge (Supplementary Fig.1a-b). Furthermore, while both virus-induced (Vir-CD8) and tumor-induced (Tum-CD8) cells progressively reacquired the expression of Bcl2, its level remained lower at all time points in Tum-CD8 cells (Fig.1d).

We then compared the phenotype of memory CD8 T cells generated following both challenges. Compared to naive CD8 T cells, Vir-CD8 and Tum-CD8 memory cells expressed surface marker associated with memory (Fig.1d-e). However, Tum-CD8 memory cells exhibited reduced expression of NKG2D, as well as the integrins CD29, CD49a, and CD49d, which are involved in the migration of antigen-induced memory CD8 T cells to inflamed tissues (33–36). Additionally, there was a lesser reduction in the expression of CD160 (Fig. 1e, Supplementary Fig. 1c). Moreover, Tum-CD8 memory cells expressed higher levels of CX3CR1, CD49b, CD49f, CD11b and, more importantly, CD43 compared to Vir-CD8 memory cells (Fig. 1e, Supplementary Fig. 1c). Notably, the expression of CD43 is inversely correlated with recall response capacity (37).

**Figure 1.**
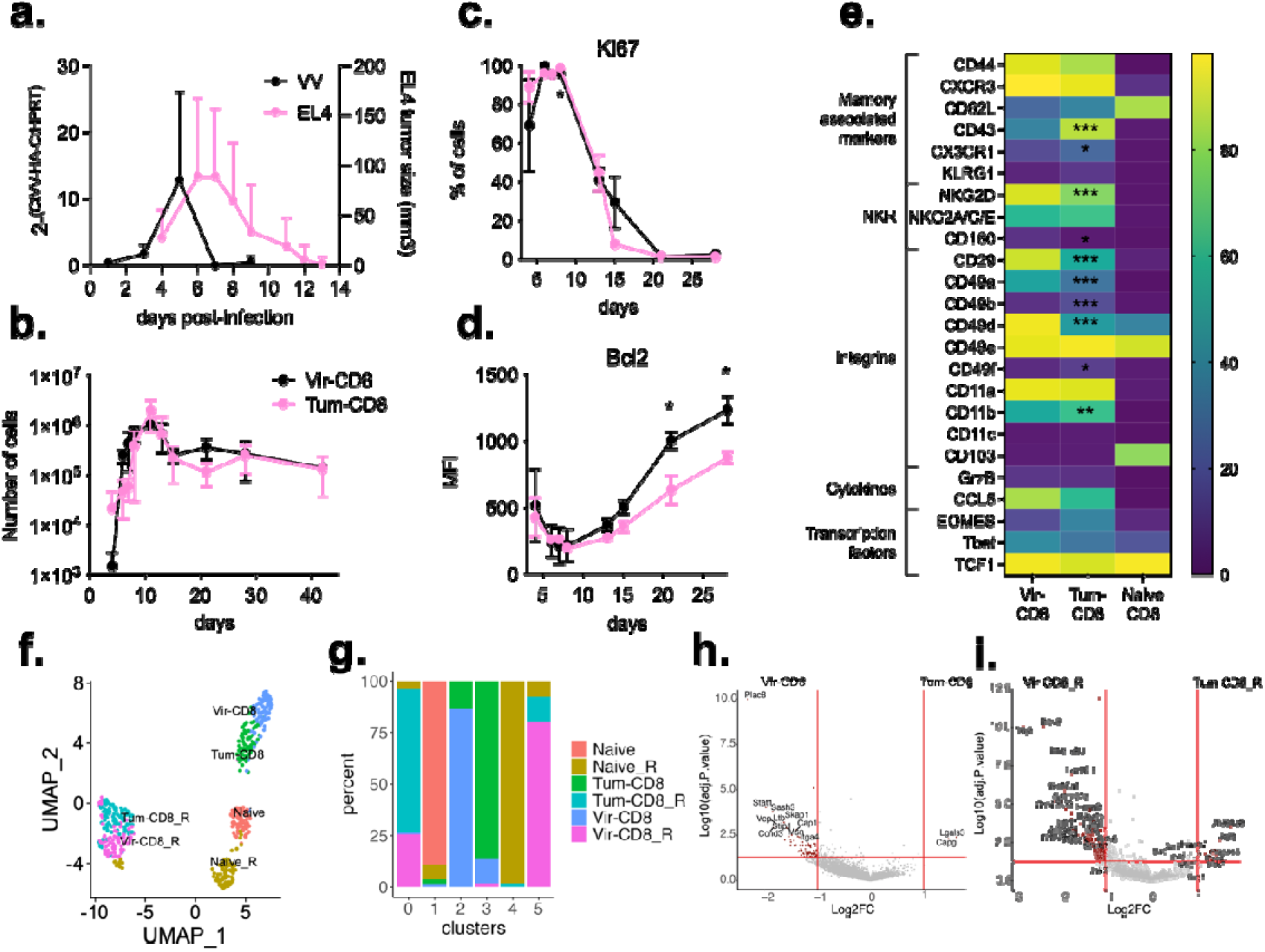
Phenotypic and transcriptional differences of memory CD8 T cells generated after a viral or a tumoral challenge. Naive F5 x CD45.1 cells (2.10^5^) were i.v. transferred in B6 mice 1-day prior immunisation with VV-NP68 (i.n., 2.10^5^ pfu) or EL4-NP68 cells (s.c., 2,5.10^6^ cells). (**a**) Viral load was measured in the lung by qPCR or tumor volume (mm^3^) was assessed by measuring its length, width and thickness over time. (**b**) The number of Vir- and Tum-CD8 cells was determined over time in the blood by flow cytometry. **(c-d**) The expression of Ki67 (**c**) and Bcl2 (**d**) by Vir- and Tum-CD8 cells was measured over time in the blood. (**e**) The phenotype of Vir- and Tum-CD8 cells was analysed 31 days after infection in the spleen and the percentages of cells expressing each marker are represented as an heatmap. The statistical significance of differences was determined using a two-way ANOVA (**c-e**) (* p <0.05, ** p <0.01, *** p <0.001). Data are represented as mean ± SD and are representative of 3 independent experiments (n= 5 to 10 mice per group). (**f-i**) 60 days after immunisation, naive F5 and Vir- and Tum-CD8 cells were single-cell sorted, and stimulated with NP68 peptide (10 nM) for 2 hours or left untreated. The transcriptome was analysed by scRNAseq (n = 476 cells). (**f**) Clustering of cells projected on a UMAP coloured by clusters. (**g**) Proportion of sorted populations in each cluster. (**h-i**) Volcano plot of the differentially expressed genes between quiescent (**h**) or restimulated (**i**) Tum-CD8 and Vir-CD8 cells.

To fully characterize Vir-CD8 and Tum-CD8 memory cells, we analyzed the whole transcriptome at the single-cell level both *ex vivo* (Vir-CD8, Tum-CD8, Naive) and following a brief *in vitro* stimulation with NP68 peptide (Vir-CD8_R, Tum-CD8_R, Naive_R). Naive CD8 T cells were included as a control. We performed a clustering analysis and projected the cells onto a UMAP (Fig.1f). We identified six clusters that were mainly defined by the experimental conditions, with memory cells generated by both challenges clustering together according to their activation status, and at a distance from naive cells (Fig.1g, Supplementary Fig.2a). This indicates that the memory cells generated following both challenges shared a significant portion of their memory gene expression signatures.

To unravel the differences in the transcriptomic profiles of Tum-CD8 and Vir-CD8 memory cells, we performed a differential expression analysis between these two cell types in quiescent (Fig.1h) or activated conditions (Fig.1i). Under quiescent condition, Tum-CD8 memory cells expressed higher levels of only two gene transcripts (*Lgals3* and *Capg*) but showed decreased levels of a larger number of genes, when compared to Vir-CD8 memory cells (Fig.1h, Supplementary Fig.2b, Supplementary Ta- bles1-2). These are associated with antiviral responses, such as *Stat1*, metabolic processes, such as *Vcp*, *Eno1* or *Gimap7* and cell adhesion, such as *Itga4*. Following *in vitro* activation, 12 genes were upregulated by Tum-CD8 memory cells compared to Vir-CD8 memory cells, including genes involved in effector responses (*Icos*, *Ccr7*, *Prf1*, *Cd9*), genes encoding inhibitory receptors (*Pdcd1*, *Havcr2*), as well as *Capg* and *Lgals3*. In contrast, Tum-CD8 memory cells did not upregulate a large number of genes related to cell metabolism and nuclear transport, as Vir-CD8 memory cells did (Fig.1i, Supplementary Fig.2b, Supplementary Tables.1-2).

Globally, memory cells generated after a transient tumor challenge exhibit decreased Bcl2 and integrin expression, along with upregulation of genes encoding inhibitory receptors.

### Tum-CD8 memory cells express molecules associated with T cell exhaustion

We showed that quiescent or activated Tum-CD8 memory cells overexpressed genes such as *Pdcd1* (PD1), *Havcr2* (TIM-3), *Lgals3* (Gal3), and *Cd9* (CD9). Increased surface expression of all these markers except Gal3 was validated at the protein level for quiescent Tum-CD8 memory cells, compared to Vir-CD8 memory cells (Fig.2a-b). Gal3 is a lectin with multiple functions in T cell biology (38). Specifically, it can alter cell functions and cytokine production by destabilizing of the secretory synapse or modifying TCR sensitivity (39–41). Secreted Gal3 can bind to glycosylated proteins such as CD44 on the surface of T cells (38) or be recruited intracellularly to the immunological synapse upon TCR stimulation (42,43). Thus, we determined the intracellular content of Gal3 and found that Tum-CD8 memory cells contained higher levels of Gal3 than did Vir-CD8 memory cells (Fig.2c-d).

**Figure 2.**
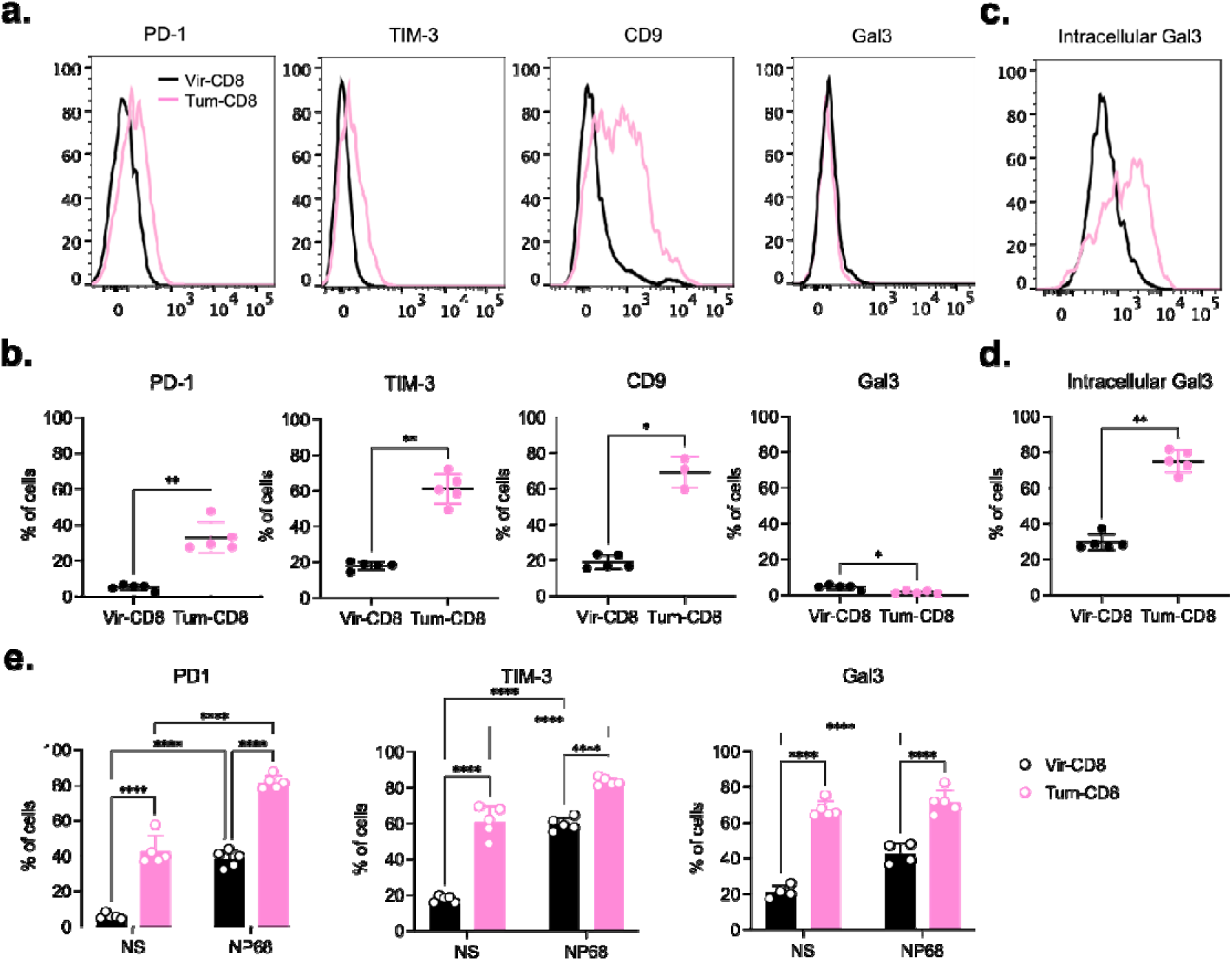
Tum-CD8 memory cells express molecules associated with T cell exhaustion. Naive F5 x CD45.1 cells (2.10^5^) were i.v. transferred in B6 mice 1-day prior immunisation with VV-NP68 (i.n., 2.10^5^ pfu) or EL4-NP68 cells (s.c., 2,5.10^6^ cells). (**a-b**) The expression of PD1, TIM-3, CD9 and Gal3 was measured at the surface of CD8 memory cells at 30 dpi by flow cytometry. (**c-d**) The expression of Gal3 was measured intracellularly in memory CD8 T cells at 30 dpi by flow cytometry. (**e**) At 30 dpi, splenocytes were stimulated with NP68 peptide (10 nM) for 4 hours, and the expression of PD1, TIM-3 or intracellular Gal3 by memory CD8 T cells was measured by flow cytometry. The statistical significance of differences was determined using Mann-Whitney test (**b; d**) or two-way ANOVA (**e**) (* p <0.05, ** p <0.01, **** p <0.0001). Data are represented as mean ± SD (n=5 mice per group) and are representative of 3 independent experiments.

The expression of these proteins was measured following NP68 peptide stimulation (Fig.2e). As observed at the genes level, the expression of both PD1 and TIM-3 increased following activation but remained higher in Tum-CD8 memory cells at all time points compared to Vir-CD8 memory cells. However, intracellular Gal3 expression remained largely unchanged following *in vitro* stimulation, particularly in Tum-CD8 memory cells. In summary, a transient tumoral challenge was sufficient to generate memory CD8 T cells that express a restricted number of exhaustion markers.

### Cytokines production but not cytotoxicity is altered in Tum-CD8 memory cells

To define the functional consequences associated with the differential expression of Gal3 and inhibitory receptors in Tum-CD8 memory cells, we analyzed their cytokine production and cytotoxic capacities. We first showed that following *in vitro* stimulation with NP68, both memory cell types exhibited comparable IL-2 production capacity (Fig.3a-b). In contrast, a smaller fraction of Tum-CD8 memory cells produced IFNγ and TNFα compared to Vir-CD8 memory cells (Fig.3a). Furthermore, based on the MFI value, the level of intracellular cytokines was significantly higher in Vir-CD8 memory cells (Fig.3b), suggesting that the cytokine production capacity of Tum-CD8 memory cells was impaired. This was further confirmed by time-course analysis of cytokine production at the cellular level and in the supernatant. Indeed, IFNγ and TNFα production was consistently lower in Tum-CD8 compared to Vir-CD8 memory cells (Fig.3c-e). Furthermore, the levels of CD69 expressed at the cell surface remained lower in Tum-CD8 memory cells than in Vir-CD8 memory cells at all time-points (Fig.3c-d).

**Figure 3.**
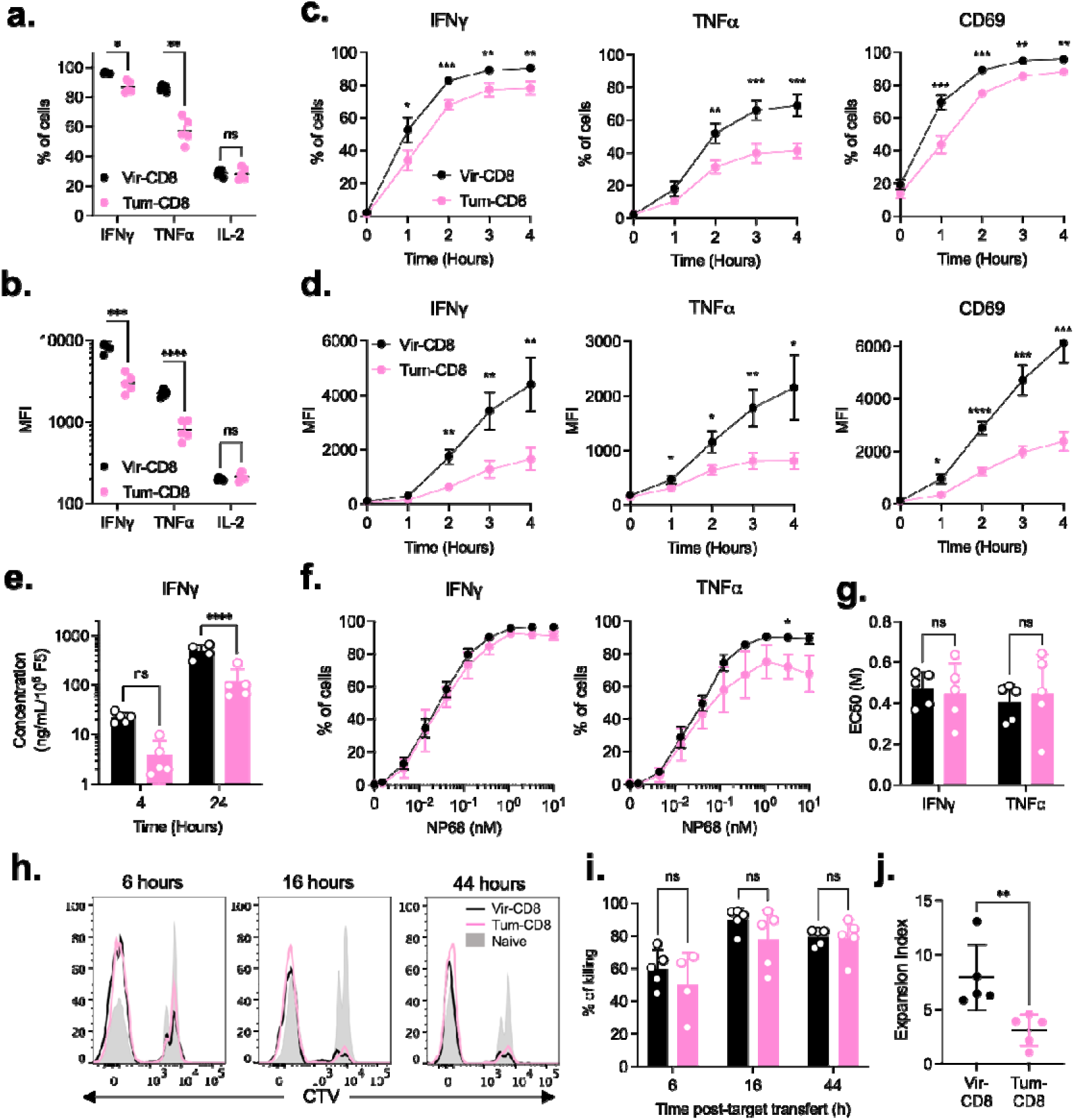
Tum-CD8 memory cells display altered cytokines production but not cytotoxic capacities compared to Vir-CD8 memory cells. Naive F5 x CD45.1 cells (2.10^5^) were i.v. transferred in B6 mice 1-day prior immunisation with VV-NP68 (i.n., 2.10^5^ pfu) or EL4-NP68 cells (s.c., 2,5.10^6^ cells). (**a-b**) At 30 dpi, F5 memory cells were restimulated with NP68 (10 nM) for 4h in the presence of GolgiStop. The production of IFNγ, TNFα and IL-2 was measured by flow cytometry and expressed in % (**a**) or MFI (**b**). (**c-d**) At 30 dpi, F5 memory cells were restimulated with NP68 (10 nM) for 4h in the presence (cytokines) or absence (CD69) of GolgiStop. The production of IFNγ and TNFα and the upregulation of CD69 were measured by flow cytometry over time and expressed in % (**c**) or MFI (**d**). (**e**) The production of IFNγ was measured in supernatant after 4 or 24h of stimulation. (**f-g**) At 30 dpi, F5 memory cells were restimulated with various doses of NP68 for 4h in the presence of GolgiStop, and the production of IFNγ and TNFα was measured by flow cytometry (**f**). EC 50 was determined (**g**). (**h-i**) Splenocytes were incubated with NP68 (10 nM) or control medium for 2h and labelled with CTV or CFSE respectively. A 1:1 ratio of NP68-loaded splenocytes: control splenocytes (2.10^6^ cells) was injected i.v. in Tum- or Vir-CD8 challenged mice at the memory stage. Representative histograms depicting control and CTV-labelled NP68-loaded splenocytes are shown (**h**). The percentage of NP68-loaded splenocytes killed was evaluated at 6-, 16-, or 44-hours post-transfer (**i**). (**j**)Total CD8 enriched from Tum- or Vir-CD8 challenged mice were labelled with CTV and stimulated with NP68-loaded DCs (1:1 ratio) for 4 days in the presence of IL-2. The expansion index of F5 cells was determined after 4 days. The statistical significance of differences was determined using 2-two ANOVA (* p <0.05, ** p <0.01, *** p <0.001, *** p <0.0001). Data are represented as mean ± SD (n= 5 mice per group) and are representative of 3 independents (**a-g**) or 1 (**h-j**) experiment(s).

As the expression of Gal3 has been associated with decreased antigen sensitivity of CD8 T cells following chronic infection (40), the antigen sensitivity of memory CD8 T cells was assessed. Cells were stimulated with increasing doses of the NP68 peptide and their cytokines production was measured after 4 h. Although the frequency of Tum-CD8 memory cells producing IFNγ- and TFNα was lower at most peptide concentrations (Fig.3f), the NP68 concentration required to achieve 50% of the maximum cytokines production (effective concentration 50 [EC50]) was similar for Tum-CD8 and Vir-CD8 memory cells (Fig.3g, Supplementary Fig.3).

Next, we evaluated the cytotoxic function of the memory CD8 T cells *in vivo*. Thirty days after the viral or tumoral challenge, mice were injected with a 1:1 ratio of NP68- loaded splenocytes to control splenocytes and the killing of NP68-loaded splenocytes was assessed at various times post-transfer. A similar frequency of NP68-loaded cells was eliminated in both groups of mice at all-time points, indicating equivalent cytotoxic activity between Tum-CD8 and Vir-CD8 memory cells (Fig.3h-i).

Finally, we measured the proliferative capacity of memory CD8 T cells in response to an *in vitro* stimulation with NP68-loaded bone marrow dendritic cells (BMDC). After 4 days of culture, living CD8 T cells were counted and the expansion index was measured. The proliferative capacity of Tum-CD8 memory cells was lower than that of Vir-CD8 memory cells (Fig.3j). Overall, the memory cells generated following a transient tumoral challenge exhibited a reduced capacity for proliferation and production of TNFα and IFNγ following secondary *in vitro* activation. However, their cytotoxic activity remained unaffected.

### A transient tumoral challenge is sufficient to impair the protective capacity of F5 memory cells

Given that both virus and tumor cells were eliminated during the challenge, we next tested whether CD8 memory cells were able to control a subsequent viral infection. Mice challenged with either virus or tumor were infected with VV-NP68 (intranasally) 30 days after the primary immunization. Six days later, the number and phenotype of F5 CD8 cells were analyzed in the lung tissue and vasculature. The proportion of Tum-CD8 memory cells within the lung tissue was lower than that of Vir-CD8 memory cells following viral challenge (Fig.4a). To assess the capacity of memory CD8 to be recruited to the lung, we determined the total number of F5 memory cells in the spleen and lung (Supp.Fig4) and calculated the fraction of these cells recruited to the lung tissue, for each mouse. Although the proportion of Tum-CD8 and Vir-CD8 memory cells within the lung tissue was similar under steady-state conditions, there was a strong recruitment of Vir-CD8 memory cells, but not Tum-CD8 memory cells or naive cells, to the lung tissue upon viral challenge (Fig.4b). This indicated that Vir-CD8 memory cells have a better capacity to migrate to the infected site. This was correlated with the increased expression of CD49a on Vir-CD8 T cells, but not Tum-CD8 memory cells in the lung, an integrin involved in targeting cells to the lung(33–36)(Fig.4c). However, the expression of CD49d did not differ between cells located in the vasculature and those in the tissue (Fig.4d).

**Figure 4.**
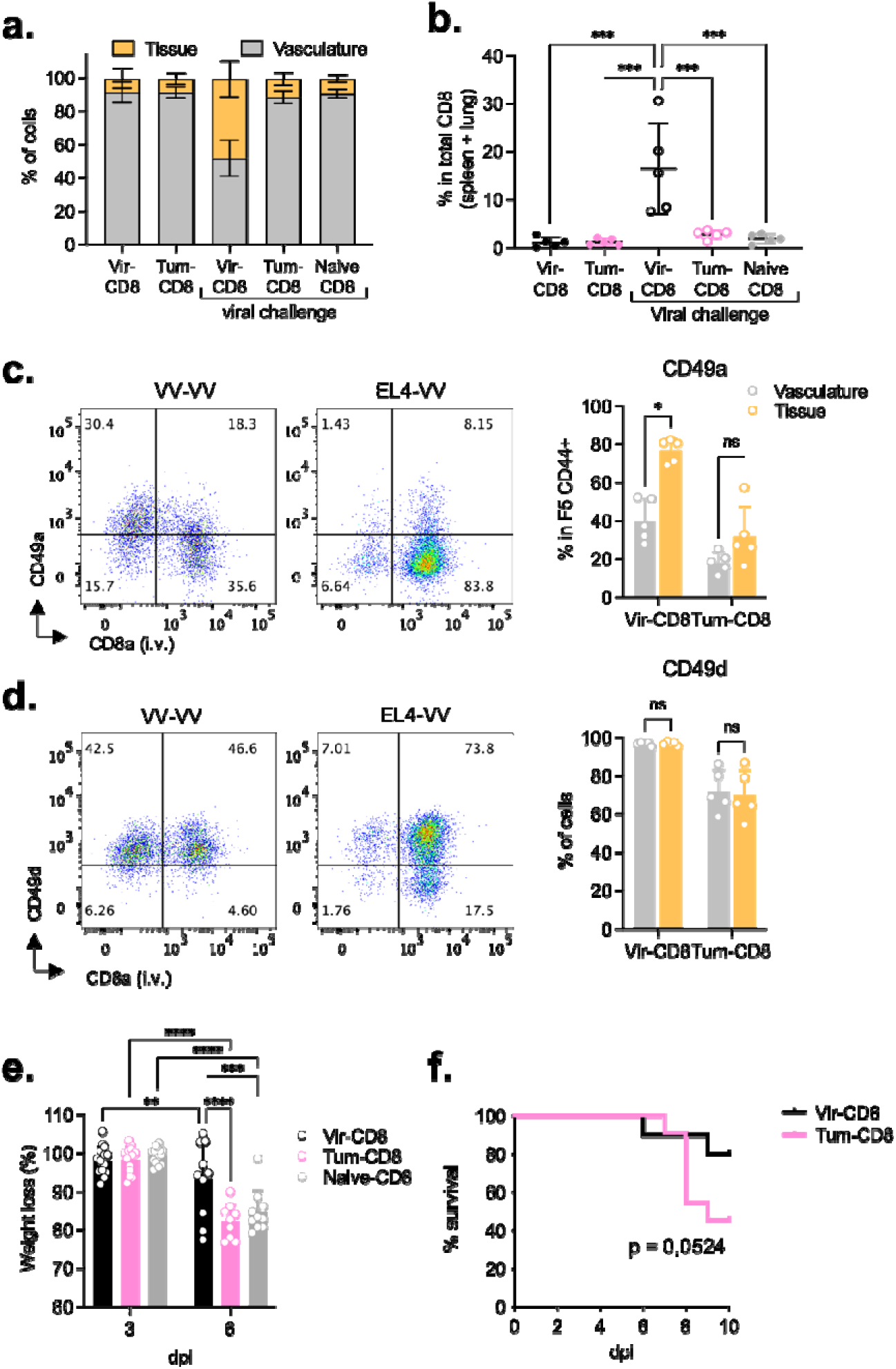
A transient tumoral challenge is sufficient to alter the protection capacity of F5 memory cells. Naive F5 x CD45.1 cells (2.10^5^) were i.v. transferred in B6 mice 1-day prior immunisation with VV-NP68 (i.n., 2.10^5^ pfu) or EL4-NP68 cells (s.c., 2,5.10^6^ cells). (**a-d**) At 30 dpi, Vir- or Tum-challenged mice were infected with VV-NP68 (2.10^5^ pfu). Six days after infection, mice received an i.v. injection of anti-CD8 antibody and the proportion of cells within the tissue and the vasculature of the lung was determined **(a**). The proportion of memory CD8 T cells in the lung tissue among all memory CD8 T cells was determined (**b**). The expression of CD49a (**c**) and CD49d (**d**) was measured on memory CD8 T cells within the lung tissue and vasculature. (**e**) At 30 dpi, Vir- or Tum-challenged mice were infected with Flu-NP68 (5.10^4^ TCID50) and the weight loss was followed for 6 days. (**f**) At 30 dpi, Vir-CD8 and Tum-CD8 memory cells were FACS-sorted and transferred into B6 host (1,2.10^5^ cells, i.v. route). One day after transfer, mice received a lethal dose of Flu-NP68 (2.10^6^ TCID 50) and survival was followed for 10 days. The statistical significance of differences was determined with 1-way (b) or 2-way (c-e) ANOVA test (* p> 0.05, ** p <0.01, *** p <0.001, **** p <0.0001), or log-rank test (**f**). Data are represented as mean ± SD (n = 5 mice per group) and are representative of 2 **(a-d, g**) or 1 (**e-f**) experiment(s).

We then wondered whether the reduced ability of Tum-CD8 memory cells to enter the lung tissue would impact their protection capacity against heterologous infection with a Flu-virus carrying the NP68 epitope. We first addressed the capacity of Tum-CD8 and Vir-CD8 memory cells to protect mice from Flu-NP68 infection in a recall context. Virus- or tumor-challenged mice were infected with Flu-NP68 (i.n.) 30 days after the first immunization. Naive mice were used as controls. Similar to naive mice, the tumor-challenged mice lost nearly 20% of their initial body weight following Flu-NP68 infection while virus-challenged mice lost only about 5% (Fig.4e).

Finally, to evaluate the role of Vir-CD8 memory cells in the protection observed upon secondary viral challenge, we sorted Tum-CD8 and Vir-CD8 memory cells at 30 dpc and transferred them into naive B6 hosts. The day after, the mice received a lethal dose of Flu-NP68, and survival rates were monitored for 10 days (Fig.4g). Mice that received Tum-CD8 memory cells exhibited a sharp decline in survival at 8 dpc, dropping to 30% survival by 10 dpc, whereas mice that received Vir-CD8 memory cells maintained an 80% survival rate, indicating superior protection (Fig. 4f). In conclusion, we demonstrated that while Tum-CD8 memory cells were capable of rejecting the primary tumor and retaining certain effector functions, they proved ineffective against viral infection.

### Phenotype and cytokines production capacity of F5 memory cells is conserved after homologous or heterologous recall

Finally, we sought to evaluate the stability of the phenotype that was imprinted after a viral or a tumoral challenge. For this purpose, viral- or tumor-challenged mice received a second homologous or heterologous recall with either VV-NP68 or EL4- NP68 at 26 dpc (Fig.5a). Mice that did not receive a second immunization were used as controls. An equivalent number of F5 memory cells was recovered from mice in all groups at 31 days post-recall (Fig.5b). The expression of the main markers that differed between Tum-CD8 and Vir-CD8 memory cells after a primary challenge was determined in the spleen memory cells. The expression of CD43 was slightly increased in Vir-CD8 memory cells following a viral recall and slightly reduced in Tum-CD8 memory cells following a recall with either a virus or a tumor (Fig.5c). However, the phenotype of Tum-CD8 and Vir-CD8 memory cells were globally stable. Indeed, the increased expression of CD9 and CD43 in Tum-CD8 memory cells was not reversed by viral challenge (Fig.5c). Furthermore, these cells did not show significantly increased expression of CD49a and CD49d, which was acquired by Vir-CD8 memory cells following a primary viral challenge (Fig.5c). Similarly, Vir-CD8 memory cells did not modify their primary phenotype following a tumor challenge (Fig.5c). The stability of the phenotype acquired during the primary challenge was also observed for cytokine production (Fig.5d) and for the expression of PD1 and TIM-3 (Fig.5e), following peptide activation.

**Figure 5.**
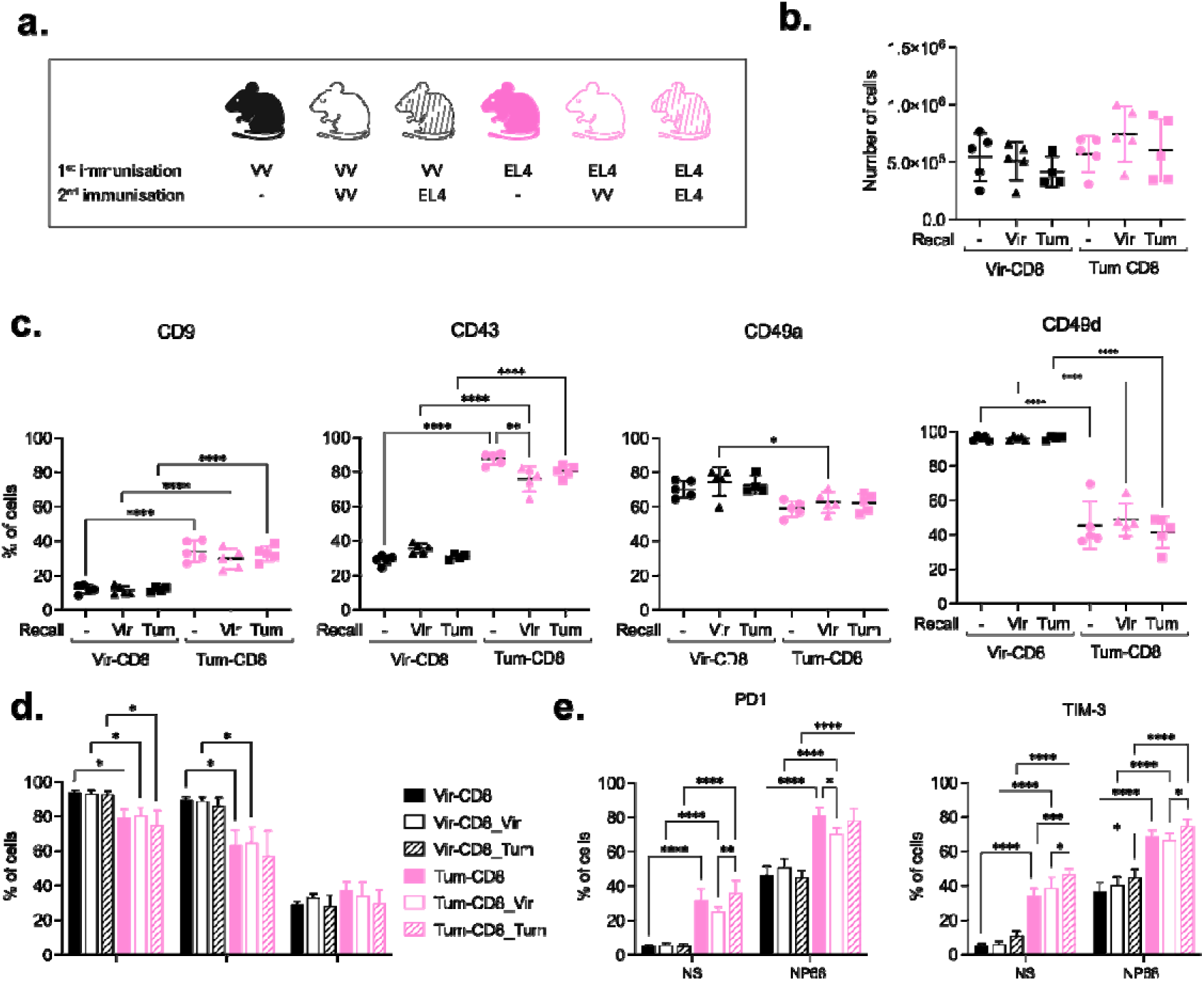
Phenotype and cytokines production capacity of F5 memory cells is conserved after homologous or heterologous recall. (**a**) Naive F5 x CD45.1 cells (2.10^5^) were i.v. transferred in B6 mice 1-day prior immunisation with VV-NP68 (i.n., 2.10^5^ pfu) or EL4-NP68 cells (s.c., 2,5.10^6^ cells). At 26 dpi, mice received a second immunisation with VV-NP68 or EL4-NP68. (**b**) Thirty-one days post-recall, the number of F5 cells was measured in the spleen. (**c**) The expression of CD9, CD43, CD49a and CD49d was measured on F5 memory cells 31 days after recall by flow cytometry. (**d-e**) Splenocytes were stimulated with NP68 (10 nM) for 4h in the presence (**d**) or absence (**e**) of GolgiStop. (**d**) The production of IFNγ, TFNα and IL-2 was measured by flow cytometry. (**e**) The expression of PD1 and TIM3 on F5 memory cells was determined in the spleen. The statistical significance of differences was determined with 1-way (b-c) or 2-way (d-e) ANOVA test (* p> 0.05, *** p <0.001, **** p <0.0001). Data are represented as mean ± SD (n = 5 mice per group) and are representative of 2 independent experiments.

Overall, a transient viral or tumoral challenge is sufficient to imprint a stable phenotype that affects the quality of memory cells.

## Discussion

In this study, we addressed the impact of a transient tumor challenge on the generation of memory CD8 cells. While the generation of memory CD8 cells was not altered by a tumoral challenge, we showed that it was sufficient to alter their quality and imprint an exhausted-like phenotype, defined by the expression of inhibitory receptors and altered cytokine production and recall responses.

Quantitatively, we showed that tumor and virus immunization led to the same number of CD8 memory cells. Similarly, it has been shown that CD8 T cells exhibit comparable expansion during the first days following transfer in mice with established liver tumors or during acute infections (31). Qualitatively, single-cell RNA-seq analyses revealed similar transcriptomic profiles in Vir-CD8 and Tum-CD8 memory cells, and differential gene expression analysis highlighted differences in the level of gene expression rather than distinct gene expression patterns between Vir-CD8 and Tum-CD8 memory cells. However, the limited number of cells analyzed (approximately 80 per experimental condition) may not have been sufficient to capture subtle differences in the gene expression profiles.

However, we demonstrated that Tum-CD8 memory cells were characterized by increased expression of several markers implicated in the restriction of T cell functions, such as Lgals3 (Gal3), Cd9 (CD9), Pdcd1 (PD1), and Havcr2 (TIM-3) genes, both at the RNA and protein levels. Some of these markers contribute to T cells exhaustion. Specifically, PD1 and TIM-3 are two inhibitory receptors whose sustained expression is a hallmark of exhausted T cells (15). PD1 restricts the expression of cytokine, such as IFNγ or TNFα, in an IRF4-dependent manner (44) while TIM-3 alters IL-2 production and limits the upregulation of CD69 (45). Gal3 also plays a role in inducing the exhaustion program, possibly through excessive activation of NFAT signaling (41). Notably, Gal3-deficient CD8+ T cells displayed increased IFNγ and IL-2 production as well as enhanced proliferation compared to WT CD8 T cells (41,43). Additionally, Gal3 and CD9 are negative regulators of immunological synapse (42,43,46). Gal3 can co-localize with Zap70 and restrict TCR signaling (43), whereas CD9 competes with ICAM-1 for LFA-1 adhesion (46).

We demonstrated that the exhausted-like phenotype exhibited by Tum-CD8 memory cells is associated with altered functionality. Specifically, these cells showed a reduced capacity to upregulate CD69 and produce IFNγ and TNFα cytokines in response to *in vitro* TCR stimulation. Furthermore, Tum-CD8 T cells displayed reduced expansion capacity in response to *in vitro* stimulation compared to those generated following viral infection. However, Tum-CD8 memory cells retained their capacity to kill target cells *in vivo* and to produce IL-2 following peptide restimulation. This finding contrasts with studies showing that IL-2 production is the first function lost during the exhaustion process in the context of chronic infections with LCMV (15,47).

In addition to their expression of inhibitory receptors, Tum-CD8 memory cells were characterized by increased expression of the activation-associated glycoform of CD43, whose expression is associated with a lower recall capacity (37). In agreement with these findings, we showed that the capacity of Tum-CD8 memory cells to protect against a Flu-infection was impaired as compared to Vir-CD8 memory cells. Furthermore, Tum-CD8 memory cells showed decreased expression of the integrins CD49a and CD49d. CD49a upregulation by CD8 T cells is driven by inflammatory cytokines such as TGFβ, IL-6, and IL-12 (48). Thus, the lower CD49a expression observed in Tum-CD8 memory cells compared to Vir-CD8 memory cells may reflect the lack of an inflammatory environment during tumor challenge. Notably, both CD49a and CD49d integrins are crucial for lung tissue homing (33–36). Consistent with their phenotype, we demonstrated that Tum-CD8 memory cells were significantly less efficient than Vir-CD8 memory cells in infiltrating lung tissue following recall with VV-NP68.

Exhausted CD8 T cells arising in chronic infections and cancer exhibit environment-specific characteristics (18,22,25). For instance, NFAT5 regulates T cell exhaustion specifically in the hyperosmotic tumor microenvironment but not in chronic viral infections (49). Here, we demonstrated that CD8 T cell priming by a tumor is sufficient to impair memory CD8 T cell functionality and promote an exhausted-like phenotype, even in the absence of tumor growth. Furthermore, we showed that CD8 T cells activated in a tumoral context exhibited defects in inflammation-associated genes expression such as *Stat1*, compared to those generated following viral infection. These findings highlight the crucial role of inflammatory signals in the generation of functional memory CD8 T cells.

The exhausted program imprinted following chronic infection is stable and cannot be reversed upon PD1 blockage (50). Similarly, we demonstrate here that the exhausted like-phenotype imprinted upon a transient tumoral challenge is stable and irreversible, as Tum-CD8 memory cells retained their distinct phenotypic traits and defective cytokine production upon both homologous and heterologous recall with VV-NP68. This aligns with recent findings showing that a tumoral challenge induces epigenetic modifications at the *Pdcd1* locus within the first hours following CD8 T cell activation (31). Further studies are needed to determine the epigenetic changes associated with a transient tumoral challenge. Similarly, Vir-CD8 memory cells maintained their phenotype and function upon both homologous and heterologous recall, indicating that the positive traits imprinted in virus-induced memory cells were stable and could not be reversed by the suboptimal-stimulation associated with the tumor.

Chronic infections are classical models used to study exhaustion. Our study emphasizes the importance of investigating the mechanisms driving CD8 T cell dysfunction associated with suboptimal-stimulation.

## Material and methods

### Mice

B6 (C57BL/6J) mice were purchased from Charles River Laboratories. F5 TCR-transgenic mice on a B6 background (B6/J-Tg(CD2-TcraF5,CD2-TcrbF5)1Kio/Jmar) (51) were crossed with B6-Ptprc^em(K302E)Jmar^/J) (52) to obtain F5 x CD45.1 mice (B6- ^Ptprcem(K302E)Jmar^/J-Tg(CD2-TcraF5,CD2-TcrbF5)1Kio/Jmar). Mice were bred and housed under specific pathogen-free conditions in the AniRA-PBES animal facility (Lyon, France). All experiments were approved by our local ethics committee (CECCAPP, Lyon, France), and accreditations were obtained from governmental agencies.

### Immunizations

The recombinant vaccinia virus (VV) expressing the NP68 epitope (VV-NP68) was engineered from VV (strain Western Reserve) by Dr. Denise Yu-Lin Teoh in Pr. Sir Andrew McMichael’s laboratory at the MRC (Human Immunology Unit, Institute of Molecular Medicine, Oxford, UK). The recombinant influenza virus strain WSN encoding the NP68 epitope (Flu-NP68) was produced by reverse genetics from the A/WSN/33 H1N1 strain (51). EL4 lymphoma cell line expressing the NP68 epitope (EL4-NP68) was provided by Dr. T.N.M. Schumacher. Naive F5 x CD45.1 cells (2.10^5^under 200 μL) were transferred by intravenous (i.v.) injection into B6 hosts. The next day, recipient mice were inoculated intranasally (i.n.) with VV-NP68 (2.10^5^ pfu under 20 μL) or subcutaneously (s.c.) with EL4-NP68 cells (2.,5.10^6^ under 200 μL). For recall experiments, more than 30 days post-challenge (dpc), mice received a second immunization with either VV-NP68 or EL4-NP68, similar to the primary challenge. For protection experiments mice were infected i.n. with a lethal dose of Flu-NP68 virus (2.10^6^ TCID50 under 20 μL). The fraction of surviving animals was measured daily for 10 days, mice that lost more than 20% of their initial body weight were euthanized

### Sample collection and flow cytometry analysis

Mice were bled at intervals of at least 7 days or euthanized for organ collection. In some experiments, for the detection of circulating cells, mice received an i.v. injection of an anti-CD8a-BUV395 antibody (1 μg under 200 μL, BD Biosciences), 3 minutes prior euthanasia. Spleens were mechanically disrupted and filtered through a sterile 100 μm nylon mesh filter. Lungs were dissociated using the Lung Dissociation kit (Miltenyi Biotech) and a gentleMACS™ Dissociator, according to manufacturer’s instructions. Blood volume was precisely measured and blood cells were enumerated using FlowCount fluorospheres (Beckman Coulter).

Single cell suspensions were first incubated with efluor780-coupled Fixable Viability Dye (ThermoFischer Scientific or TFS) for 20 min at 4 °C. Nonspecific binding was then blocked with Fc-blocking antibody (Ab) 2.4G2 for 10 min at 4 °C. Surface staining was performed with an appropriate mixture of Ab diluted in staining buffer (PBS supplemented with 1% fetal calf serum (Life Technologies) and 0.09% NaN3 (Sigma-Aldrich) for 30 min at 4 °C. For intracellular staining, cells were fixed and permeabilized, according to the manufacturer’s instructions, using either CytoFix/CytoPerm buffer (BD Biosciences) for cytokines staining or Fixation/Permeabilization buffer from the Foxp3 Transcription Factor Staining Buffer Kit (eBioscience) for transcription factors staining. Fixed cells were then stained with an appropriate mixture of intracellular Ab for 30 min at 4 °C. All analyses were performed using a BD LSRFortessa cell analyzer (BD Biosciences) and further analyzed using FlowJoTM v10 software (BD Biosciences). Antibodies are listed in the Supplementary Table.3.

### Cell Culture

The EL4-NP68 cells line was maintained for a maximum of 2 weeks at 37 °C in a 5% CO2 incubator and cultured in complete RPMI medium, consisting of RPMI 1640 medium with GlutaMAX™ Supplement (TFS) supplemented with 10% Fetal Calf Serum (BioWest), 50 μg/mL gentamicin, 10 mM HEPES buffer and 50 μM 2- Mercaptoethanol (50 μM) (all from TFS). Murine primary cells were cultured in DMEM medium consisting with 4.5 μg/mL of glucose and GlutaMAX™ supplement (TFS), supplemented with 6% Fetal Calf Serum (BioWest), MEM Non-Essential Amino Acids Solution, 50 μg/mL gentamicin, 10 mM HEPES buffer and 50 μM 2-Mercaptoethanol (50 μM) (all from TFS), and maintained at 37 °C in 7% CO2 incubator.

### In vitro CD8 peptide-stimulation

Splenocytes were stimulated with 10 nM NP68 peptide (ASNENMDAM, Proteogenix) for 1 to 24 h. GolgiStop (BD Biosciences) was added during the last 4 h of culture for intracellular cytokine detection. The measurement of inhibitory receptors (PD1 and TIM3) and CD69 was performed in absence of GolgiStop (BD Biosciences) as it blocks the expression of these markers at the cell surface. For the dose-response experiments, splenocytes were stimulated with NP68 peptide at a dose ranging from 10^-3^ to 10 nM, for 4 h in the presence of GolgiStop.

### In vitro proliferation assay

Peptide-loaded bone marrow-derived dendritic cells (BMDC) were used to activate CD8 T cells. Bone marrow cells were isolated by flushing femurs of B6 mice with complete RPMI medium. Cells were filtered through a 100-μm nylon cell strainer and cultured at 2.10^6^ cells/mL in complete RPMI medium supplemented with 100 ng/ml recombinant human Flt3l (Amgen). After 7 days, NP68 peptide (20 nM) and CpG ODN 1826 (2 mg/mL, InvivoGen) were added to the BMDC and cultured overnight before being washed and used for CD8 activation. Total CD8 T cells were enriched from splenocytes using an autoMACS® Pro Separator with the CD8a+ T cell isolation kit (Miltenyi Biotec) according to the manufacturer’s instructions. Enriched CD8 T cells were then labelled with CTV (CellTrace Violet, 2.5 mM, TFS) according to the manufacturer’s instructions. CTV-labelled CD8 T cells and NP68-loaded BMDC were co-cultured in a 1:1 ratio in the presence of murine 5% rIL-2 supernatant (corresponding to a final concentration of 11.5 ng/mL) for 4 days. Cells were then stimulated with NP68 (10 nM) for 4 hours in the presence of GolgiStop. The number of cells was determined using FlowCount fluorospheres (Beckman Coulter). The expansion index was calculated as follows: Number of F5 CD8 at the end of the culture/number of F5 CD8 stimulated on day 0.

### In vivo cytotoxicity assay

Total splenocytes were incubated in medium with or without 10 nM NP68 for 2h and labelled with respectively CTV or CFSE (TFS). A 1:1 ratio of NP68-loaded splenocytes: control splenocytes (2.10^6^ cells) was injected i.v. into naive or challenged mice. After 6, 16 or 44h, spleens were collected and the ratio of NP68-loaded splenocytes/control splenocytes was evaluated by flow cytometry for naive or challenged mice. The percentage of *in vivo* killing in individual challenged mice was calculated using the mean ratio of NP68-loaded splenocytes/control splenocytes in naive mice, as follows:

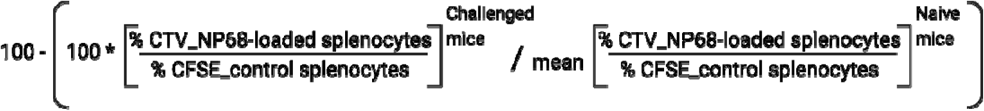

### ELISA

The concentration of IFNγ in the supernatant was quantified using ELISA MAX^TM^ Standard Set for mouse IFNγ (BioLegend) following manufacturer’s instructions. Absorbance was measured at 450 nm using an Infinite 200 microplate reader (TECAN). The IFNγ concentration was normalized to the number of F5 cells per well on day 0 and expressed in ng/mL per 10^5^ F5 cells.

### qRT-PCR

The VV-NP68 viral load was determined in the lungs after DNA extraction. The lungs were directly frozen in nitrogen and stored at −80 °C before DNA extraction. The lungs were homogenized in 150 μL of PBS using a Precellys 24 beads grinder homogenizer (Bertin Technologies) 1 for 15 s at 4 m/s. Total DNA was extracted from cell lysates using NucleoSpin Tissue kit (Macherey Nagel).

qPCR was performed in duplicate for each gene using the StepOnePlus™ Real-Time PCR System and the Platinum® SYBR® Green qPCR SuperMix-UDG with ROX. Individual data were normalized to HPRT mRNA, by calculating the ΔCt (median Ct (gene) - median Ct (HPRT)). Primers (Sigma–Aldrich) were used as follows:

VV-HA-Forward CATCATCTGGAATTGTCACTACTAAA; VV-HA-Reverse ACGGCCGACATATAATTAATGC; HPRT-Forward AAAGACTTGCTCGAGATG; HPRT-Reverse TAATGTAAT-CCAGCAGGTC.

### Single-cell RNA sequencing

CD45.1+ F5 memory cells were sorted from the spleens of VV- or EL4-immunised mice 9 weeks after the primary challenge. Naive F5 cells were sorted from the spleen of F5 x CD45.1 mice. Enriched CD8 T cells (see In vitro proliferation assay) were stimulated (Vir-CD8_R, Tum-CD8_R, naive_R) or not (Vir-CD8, Tum-CD8, Naive) with NP68 peptide (10 nM) for 2h at 37 °C. F5 cells were then single-cell sorted according to their viability and their expression of CD45.1, CD8a, and CD44, into a 96- well plate containing 4 μL of lysis solution (PBS, 0,04% Triton 0,4%, 2 U/μL RNaseOut [TFS], 2,5 μM of poly(T) reverse transcription primers and 2,5 mM dNTPs), on a FACS Aria II (BD Biosciences).

Library construction was performed following the Smart-seq2 protocol(53). Briefly, mRNA was reverse transcribed into cDNA and pre-amplified by PCR. cDNAs was then fragmented by tagmentation and Illumina sequencing adaptors containing indexes allowing for cell identification were added. The cDNA was amplified by PCR. Each 96-cells libraries were pooled together and both quality and quantity were tested using a DNA high sensitivity D1000 ScreenTape on Tapestation 4200 (Agilent). Libraries were sequenced on an Illumina HiSeq in paired-ends (2 x 150 bp) at an average sequencing depth of ∼1M reads/cells.

### scRNA-Seq analysis

Quality control of the raw data was performed using FastQC. The Nextera Transposase sequence and low-quality bases were trimmed using Trim Galore. Transcript expression was quantified using Salmon and version M23 of GENCODE mouse genome and transcriptome. The gene/count matrix was generated with tximport. Cells were filtered using the *metric_sample_filter* function implemented in SCONE package (54). For each criterion (the number of reads, the number of detected genes and the areas under the false negative rate (FNR) curve employing housekeeping genes), values greater than three median absolute deviations from the median were excluded. Moreover, genes expressed in fewer than three cells were removed.

Then, the dataset consisting of 476 cells was normalized using SCnorm (55) and variable features (HVG) were selected based on variance modeling statistics from the *modelGeneVar* function in Scran (56). The top 10% of genes with positive biological components were used for further analysis. The log-normalized expression values of the 570 HVGs were used for advanced analysis.

The top eight principal components were selected for UMAP and clustering analysis. Clustering was performed by applying the Louvain algorithm included in the Seurat package V4. The top 20 differentially expressed genes (DEG) for each group of cells were determined using the *FindAllMarkers* function (logfc.threshold=0.5 and min.diff.pct=0.25) from the Seurat package. Differential expression analysis was performed using Limma and applied to all genes and not just on HVGs (57). Heatmap, dot plot and violin plot were generated using the Seurat function *DoHeatmap*, *DotPlot* and *VlnPlot* respectively. Gene ontology (GO) enrichment analysis selecting biological process was performed using the *enrichGO* function implemented in the ClusterProfiler package. Vizualisation was made with the enrichplot package.

### Statistical analysis

Statistical analyses were performed using Graphpad software Prism 10. Groups comparison was performed using the Mann-Whitney t-test, one-away ANOVA or two-way ANOVA, as indicated.

### Data availability

The sequencing data generated in this study are available at GEO NCBI under the accession number GSE283102.

## Acknowledgment

We acknowledge the contributions of SFR BioSciences (UAR3444/CNRS, US8/Inserm, École Normale Supérieure de Lyon, Université de Lyon) and of the CELPHEDIA infrastructure (http://www.celphedia.eu/), especially the center AniRA in Lyon (AniRA-Cytométrie and AniRA-PBES facilities) and Yann Leverrier. We acknowledge Severine Valsesia and Mélanie Wencker for their help with the RNA library preparation. We acknowledge Laurent Modolo for his assistance in the analysis of single cell RNA-Seq. This work was supported by INSERM, CNRS, Université de Lyon, ENS Lyon, Région Auvergne-Rhône-Alpes (Ingerence pack ambition) and ANR (MEMOIRE ANR-18-CE45-0001). Margaux Prieux has a région Auvergne-Rhône-Alpes Ph.D. fellowship. Part of some figures were created with BioRender.com.

