## Supplementary Figures for "A temporary challenge by tumor cells can lead to a permanent partial-impairment of memory CD8 T cell function"

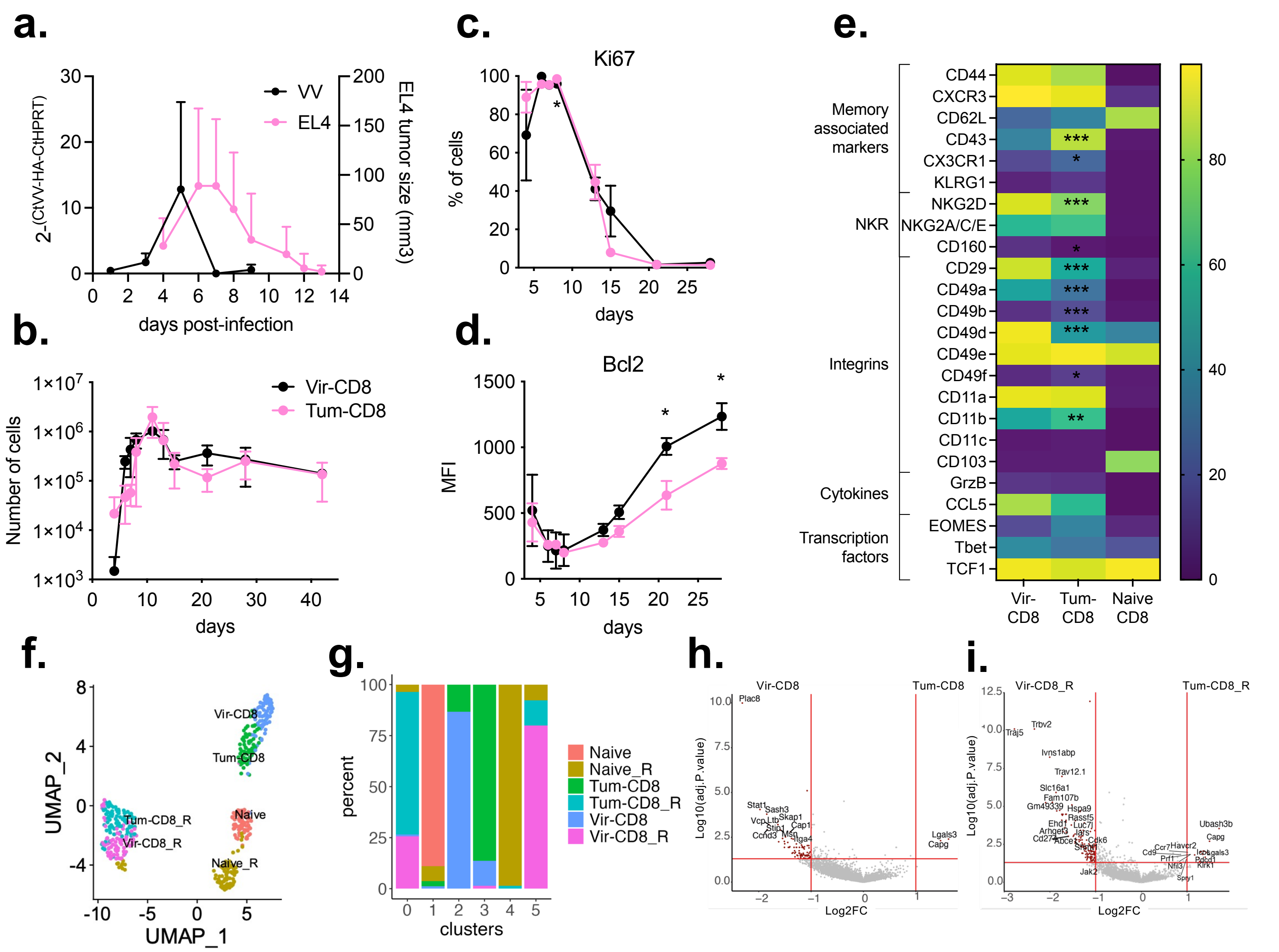


**Figure 1: Phenotypic and transcriptional differences of memory CD8 T cells generated after a viral or a tumoral challenge**

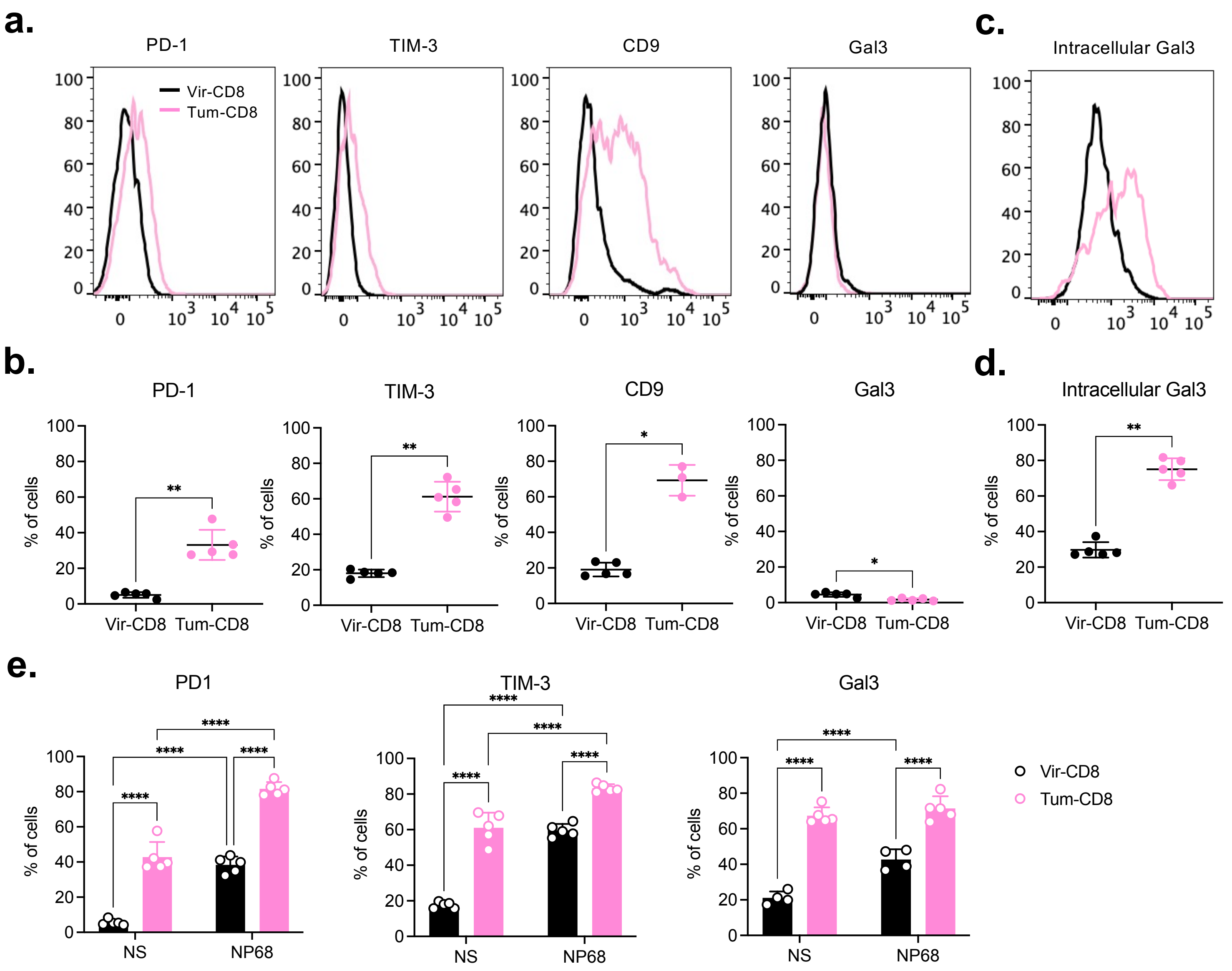


**Figure 2 : Tum-CD8 memory cells express molecules associated with T cell exhaustion**
Naive F5 x CD45.1 cells ($2.10^5$) were i.v. transferred in B6 mice 1 day prior immunisation with VV-NP68 (i.n., $2.10^5$ pfu) or EL4-NP68 cells (s.c., $2{,}5.10^6$ cells). (**a-b**) The expression of Gal3, PD1, TIM-3 and CD9 was measured at the surface of CD8 memory cells at 30 dpi by flow cytometry. (**c-d**) The expression of Gal3 was measured intracellularly in memory CD8 T cells cells at 30 dpi by flow cytometry. (**e**) At 30 dpi, splenocytes were stimulated with NP68 peptide (10 nM) for 4 hours, and the expression of PD1, TIM-3 or intracellular Gal3 by memory CD8 T cells was measured by flow cytometry. The statistical significance of differences was determined with Mann-Whitney test (* p <0.05, ** p <0.01). Data are represented as mean ± SD (n=5 mice per group) and are representative of 3 independent experiments.

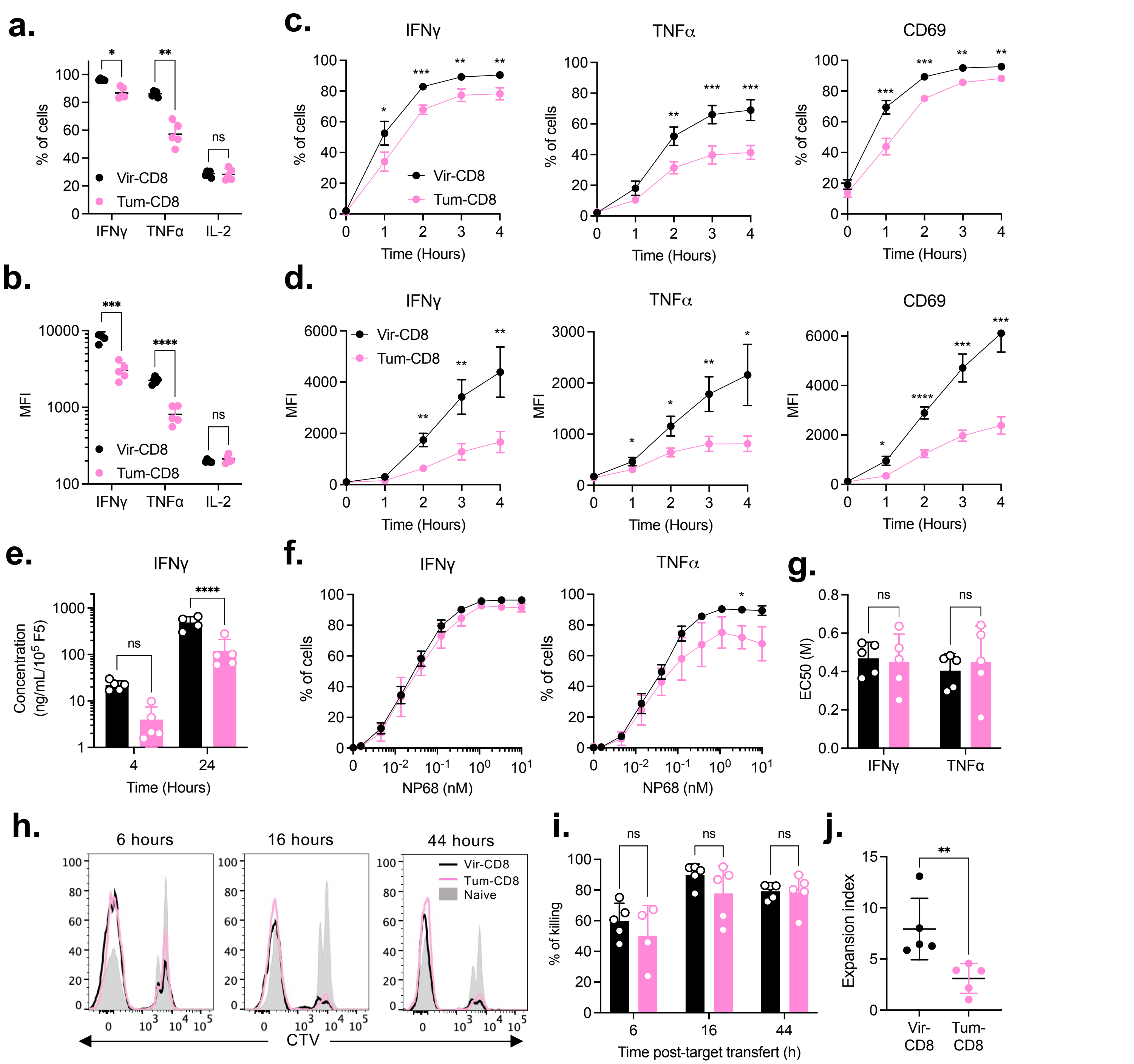

**Figure 3: Tum-CD8 memory cells display altered cytokines production but not cytotoxic capacities compared to Vir-CD8 memory cells**

Naive F5 x CD45.1 cells ($2.10^5$) were i.v. transferred in B6 mice 1 day prior immunisation with VV-NP68 (i.n., $2.10^5$ pfu) or EL4-NP68 cells (s.c., $2{,}5.10^6$ cells). (**a-b**) At 30 dpi, F5 memory cells were restimulated with NP68 (10 nM) for 4h in the presence of GolgiStop. The production of IFNγ, TNFα and IL-2 was measured by flow cytometry and expressed in % (**a**) or MFI (**b**). (**c-d**) ) At 30 dpi, F5 memory cells were restimulated with NP68 (10 nM) for 4h in the presence (cytokines) or absence (CD69) of GolgiStop. The production of IFNγ and TNFα and the upregulation of CD69 were measured by flow cytometry over time and expressed in % (**c**) or MFI (**d**). (**e**) The production of IFNγ was measured in supernatant after 4 or 24h of stimulation. (**f-g**) ) At 30 dpi, F5 memory cells were restimulated with various doses of NP68 for 4h in the presence of GolgiStop, and the production of IFNγ and TNFα was measured by flow cytometry (**f**). EC 50 was determined (**g**). (**h-i**) Splenocytes were incubated with NP68 (10 nM) or control medium for 2h and labelled with CTV or CFSE respectively. A 1:1 ratio of NP68-loaded splenocytes : control splenocytes ($2.10^6$ cells) was injected i.v. in pre-immunised mice. Representative histograms depicting the proportion of CTV-labelled NP68-loaded splenocytes are shown (**h**). The percentage of killing of NP68-loaded splenocytes was evaluated at 6, 16 ou 44 hours post-transfer (**i**). (**j**)Total CD8 were labelled with CTV and stimulated with NP68-loaded DCs (1:1 ratio) for 4 days in the presence of IL-2 and the expansion index of F5 cells was determined after 4 days. The statistical significance of differences was determined with Mann-Whitney test (** $p < 0.01$, *** $p < 0.001$). Data are represented as mean ± SD (n= 5 mice per group) and are representative of 3 independents (**a-g**) or 1 (**h-j**) experiments.

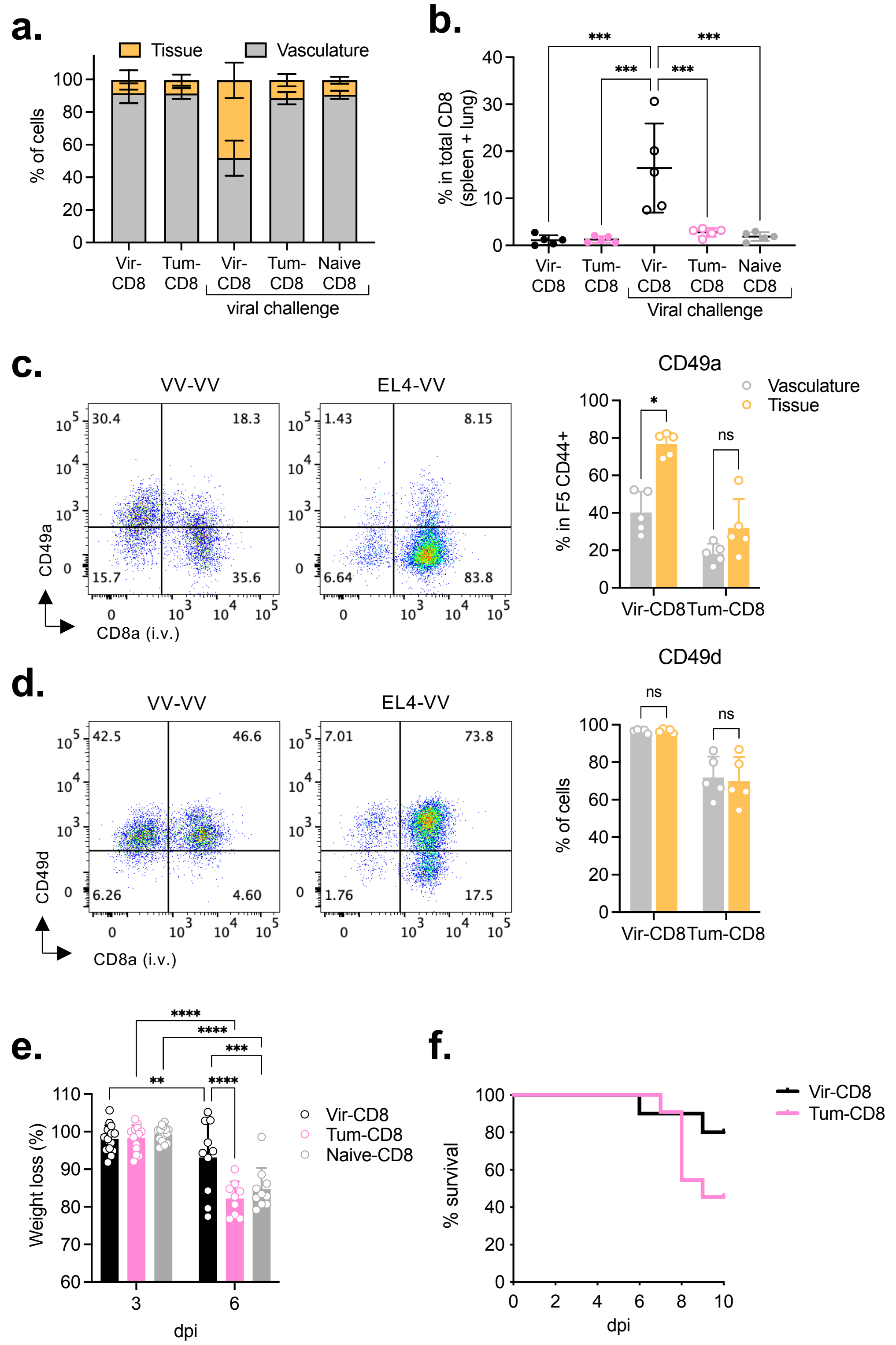


### Figure 4: A transient tumoral challenge is sufficient to alter the protection capacity of F5 memory cells

Naive F5 x CD45.1 cells (2.10⁵) were i.v. transferred in B6 mice 1 day prior immunisation with VV-NP68 (i.n., 2.10⁵ pfu) or EL4-NP68 cells (s.c., 2,5.10⁶ cells). (**a-d**) At 30 dpi, VV- or EL4-challenged mice were infected with VV-NP68 (2.10⁵ pfu). Six days infection, mice received an i.v. injection of anti-CD8 antibody and the proportion of cells within the tissue and the vasculature of the lung was determined **(a)**. The proportion of memory CD8 T cells in the lung tissue among all memory CD8 T cells was determined (**b**). The expression of CD49a (**c**) and CD49d (**d**) was measured on memory CD8 T cells within the lung tissue and vasculature. (**e**) At 30 dpi, VV- or EL4-challenged mice were infected with Flu-NP68 (5.10⁴ TCID50) and the weigh loss was followed for 6 days. (**f**) At 30 dpi, V-CD8 and T-CD8 memory cells were FACS-sorted and transferred into B6 host (1,2.10⁵ cells, i.v. route). One day after transfer, mice received a lethal dose of Flu-NP68 (2.10⁶ TCID 50) and survival was followed for 10 days. The statistical significance of differences was determined with 1-way (**b**) or 2-way (**c-e**) ANOVA test (* p> 0.05, ** p <0.01, *** p <0.001, **** p <0.0001). Data are represented as mean ± SEM (n = 5 mice per group) and are representative of 2 **(a-d, f)** or 1 (**e**) experiments.

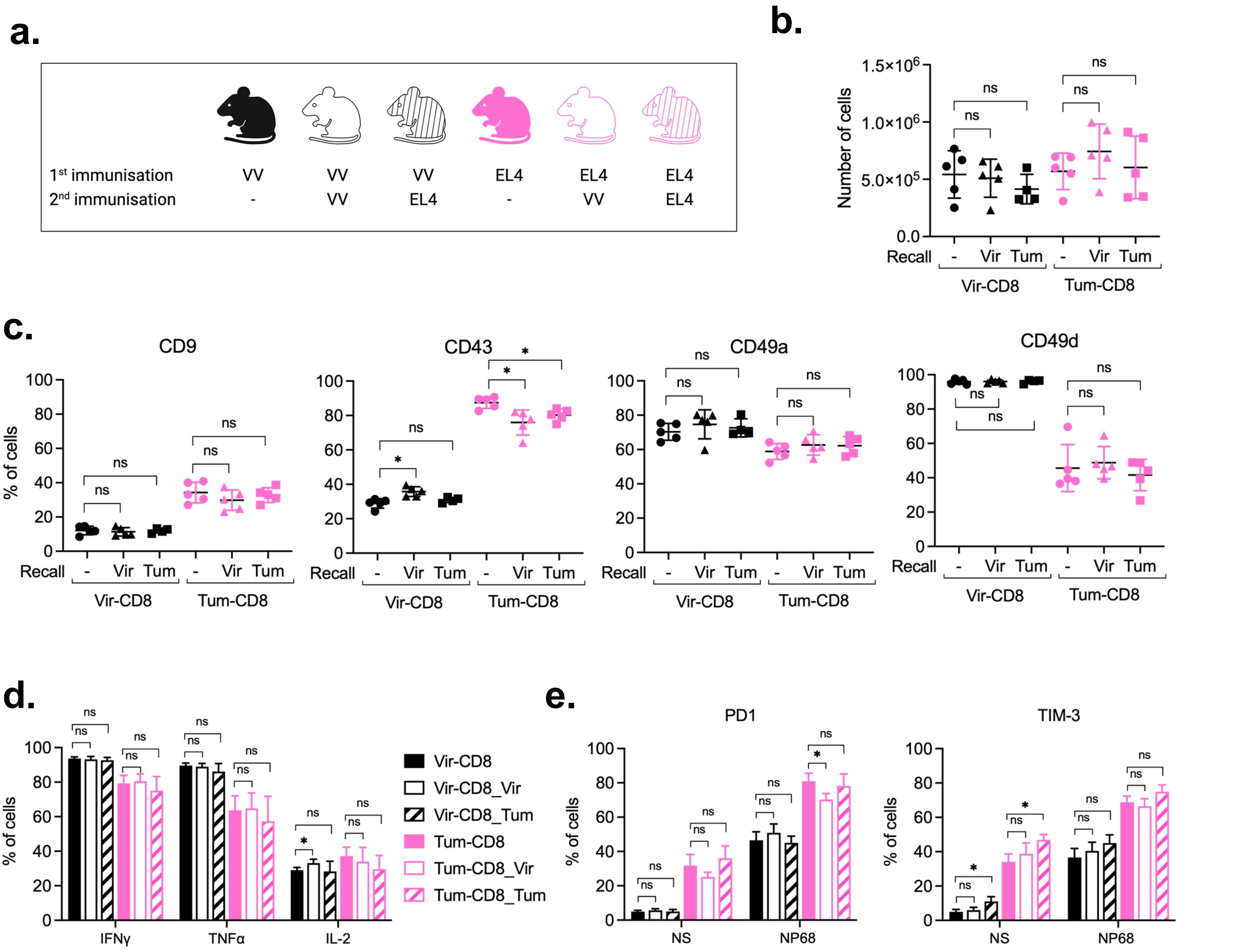


**Figure 5: Phenotype and cytokines production capacity of F5 memory cells is conserved after homologous or heterologous recall**

(**a**) Naive F5 x CD45.1 cells ($2.10^5$) were i.v. transferred in B6 mice 1 day prior immunisation with VV-NP68 (i.n., $2.10^5$ pfu) or EL4-NP68 cells (s.c., $2{,}5.10^6$ cells). At 26 dpi, mice received a second immunisation with VV-NP68 or EL4-NP68. (**b**) Thirty one days post recall, the number of F5 cells was measured in the spleen. (**c**) The expression of CD9, CD43, CD49a and CD49d was measured on F5 memory cells 31 days after recall by flow cytometry. (**d-e**) Splenocytes were stimulated with NP68 (10 nM) for 4h in the presence (**d**) or absence (**e**) of GolgiStop. (**d**) The production of IFNγ, TFNα and IL-2 was measured by flow cytometry. (**e**) The expression of PD1 and TIM3 on F5 memory cells was determined in the spleen. Statistical significance of differences was determined with Mann-Whitney test (* $p < 0.05$, ** $p < 0.01$). Data are represented as mean ± SD (n = 5 mice per group) and are representative of 2 independent experiments.

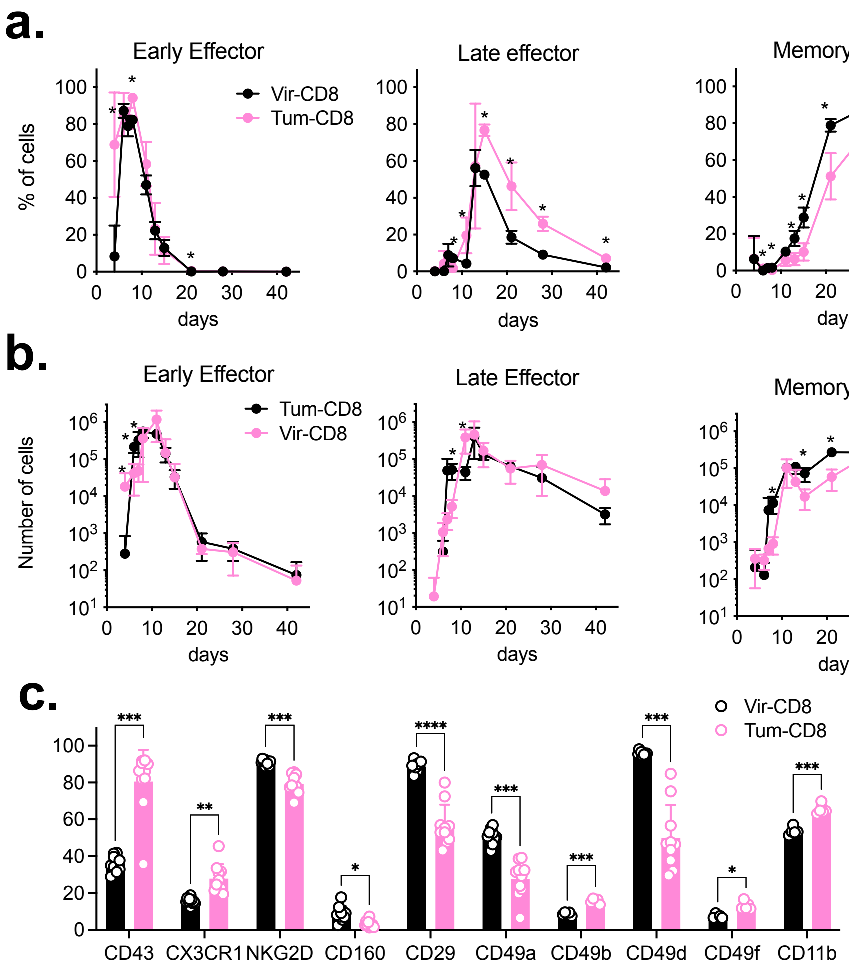


**Supplementary Figure 1: Delayed memory cells generation following transient tumoral challenge**

Naive F5 x CD45.1 cells ($2.10^5$) were i.v. transferred in B6 mice 1-day prior immunisation with VV-NP68 (i.n., $2.10^5$ pfu) or EL4-NP68 cells (s.c., $2{,}5.10^6$ cells). (**a-b**) The percentages (**a**) and the numbers (**b**) of Early effectors, Late effectors or Memory cells within Vir- and Tum-CD8 cells were determined over time in the blood by flow cytometry, using Ki67 and Bcl2 labelling. (**c**) Markers with statistical significant difference identified in Fig.1e are represented. Statistical significance of differences was determined using a two-way ANOVA (* p <0.05, ** p <0.01, *** p <0.001, **** p <0.0001). Data are represented as mean ± SD and are representative of 3 independent experiments.

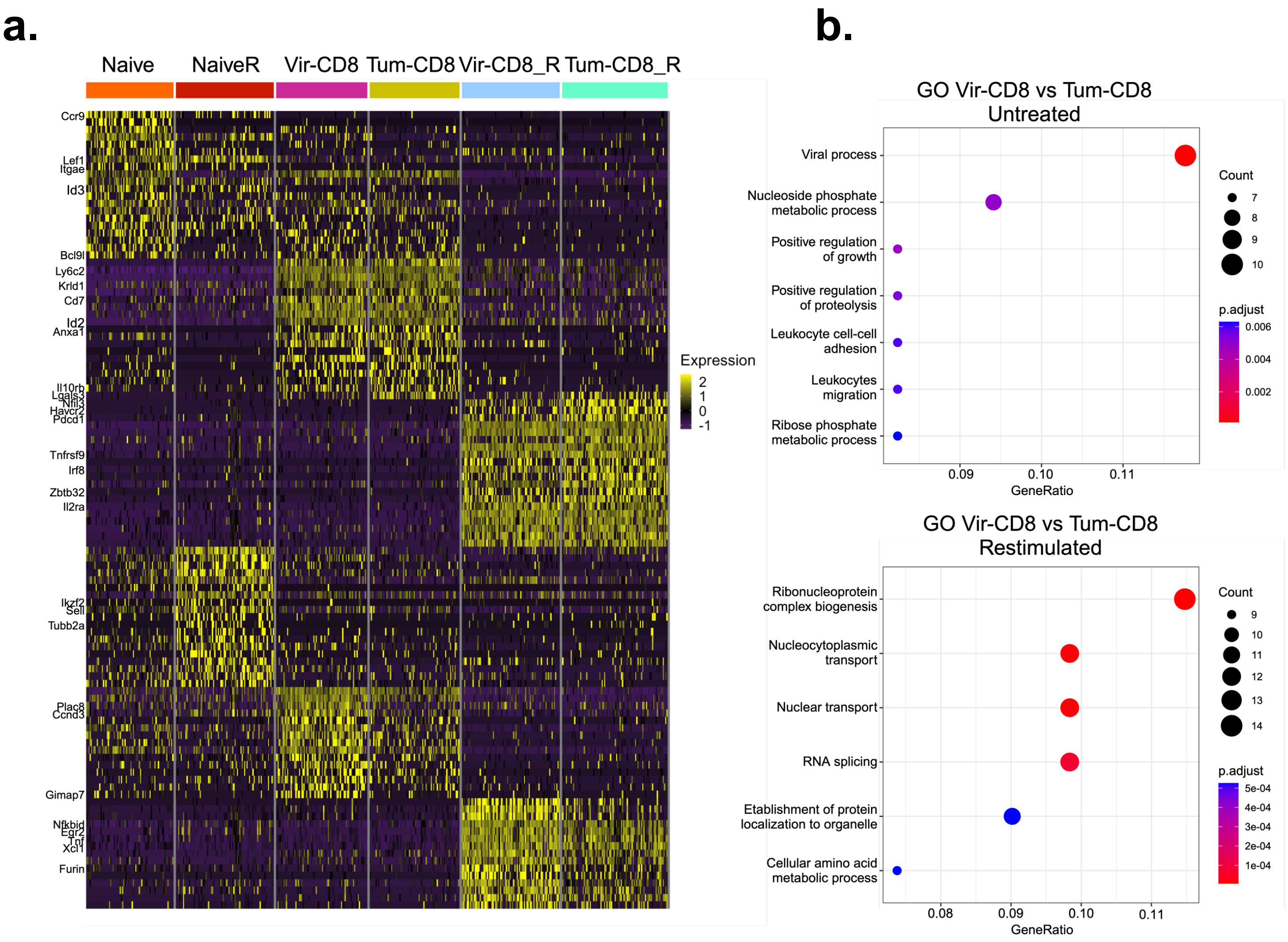


**Supplementary Figure 2: Genes differentially expressed between Tum-CD8 and Vir-CD8 memory cells**

Transcriptomic analysis of Tum-CD8 and Vir-CD8 memory cells was performed as described in Figure 1. (**a**) Heatmap of the top 20 markers for each group. (**b**) Top GO analysis of biological processes of DEGs in Vir-CD8 compared to Tum-CD8 untreated or restimulated. Node size represents the number of genes, and color intensity corresponds to the adjusted p-value.

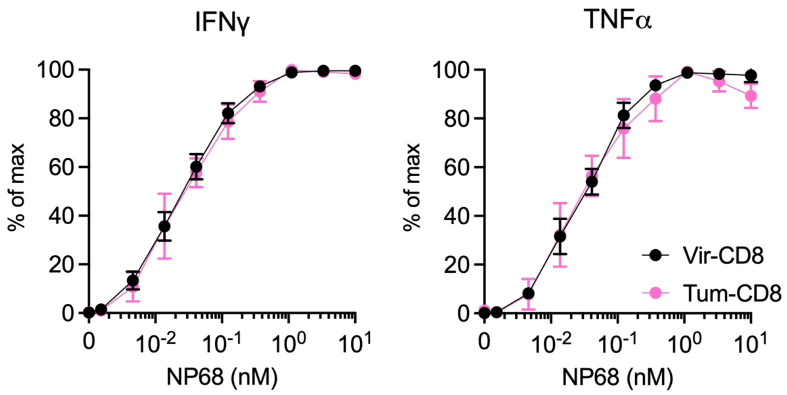


**Supplementary figure 3 : IFNγ and TNFα dose response to NP68 stimulation**

Naive F5 x CD45.1 cells ($2.10^5$) were i.v. transferred in B6 mice 1-day prior immunisation with VV-NP68 (i.n., $2.10^5$ pfu) or EL4-NP68 cells (s.c., $2,5.10^6$ cells). At 30 dpi, F5 memory cells were restimulated with various doses of NP68 for 4h in the presence of GolgiStop. The production of IFNγ and TNFα was measured by flow cytometry and expressed in percentage of maximal production. Data are represented as mean ± SD (n= 5 mice per group) and are representative of 3 independents experiments.

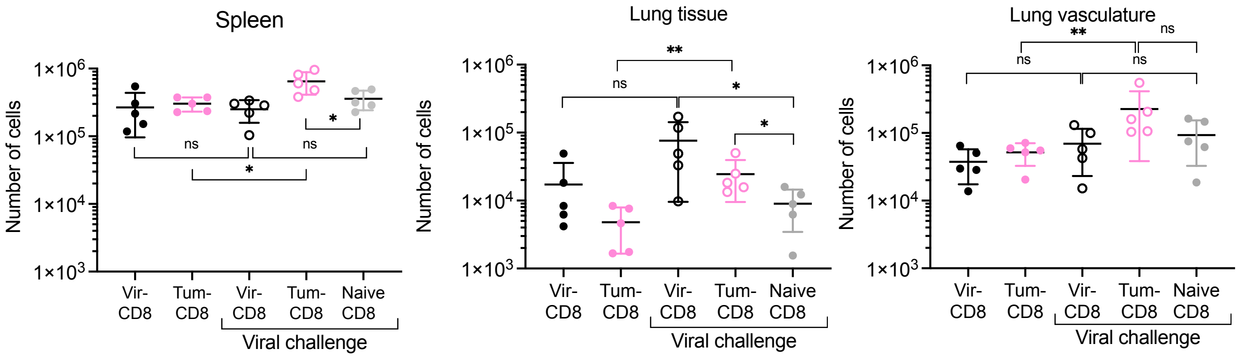


**Supp Figure 4 : A transient tumoral challenge is sufficient to alter the protection capacity of F5 memory cells**

Naive F5 x CD45.1 cells (2.10$^5$) were i.v. transferred in B6 mice 1-day prior immunisation with VV-NP68 (i.n., 2.10$^5$ pfu) or EL4-NP68 cells (s.c., 2,5.10$^6$ cells). At 30 dpi, Vir- or Tum-challenged mice were infected with VV-NP68. Six days post-challenge, mice received an i.v. injection of anti-CD8 antibody to label circulating cells. The number of CD8 memory cells in the spleen, lung tissue and lung vasculature were determined. The statistical significance of differences was determined using a one-way-ANOVA (* $p < 0.05$, ** $p < 0.01$). Data are represented as mean ± SD (n=5 mice per group).
